# Russian doll genes and complex chromosome rearrangements in *Oxytricha trifallax*

**DOI:** 10.1101/216440

**Authors:** Jasper Braun, Lukas Nabergall, Rafik Neme, Laura F. Landweber, Masahico Saito, Nataša Jonoska

## Abstract

Ciliates have two different types of nuclei per cell, with one acting as a somatic, transcriptionally active nucleus (macronucleus; abbr. MAC) and another serving as a germline nucleus (micronucleus; abbr. MIC). Furthermore, *Oxytricha trifallax* undergoes extensive genome rearrangements during sexual conjugation and post-zygotic development of daughter cells. These rearrangements are necessary because the precursor MIC loci are often both fragmented and scrambled, with respect to the corresponding MAC loci. Such genome architectures are remarkably tolerant of encrypted MIC loci, because RNA-guided processes during MAC development reorganize the gene fragments in the correct order to resemble the parental MAC sequence. Here, we describe the germline organization of several nested and highly scrambled genes in *Oxytricha trifallax*. These include cases with multiple layers of nesting, plus highly interleaved or tangled precursor loci that appear to deviate from previously described patterns. We present mathematical methods to measure the degree of nesting between precursor MIC loci, and revisit a method for a mathematical description of scrambling. After applying these methods to the chromosome rearrangement maps of *O. trifallax* we describe cases of nested arrangements with up to five layers of embedded genes, as well as the most scrambled loci in *O. trifallax*.

## INTRODUCTION

Ciliates have two types of nuclei within the same cell, where one acts as a germline nucleus (micronucleus; abbr. MIC) and the other acts as a somatic nucleus (macronucleus; abbr. MAC). The spirotrich ciliate *Oxytricha trifallax* undergoes massive genome reorganization during the post-zygotic development of its daughter cells (Prescott 1994).

Conjugation in *O. trifallax* leads to destruction of the parental, vegetative MAC and regeneration of a new MAC from a copy of the zygotic MIC. This process involves the selective removal of more than 90% of the germline genomic information, and the precise rearrangement of the remaining pieces in a particular order and orientation (Yerlici and Landweber 2014). The mature MAC contains millions of gene-sized chromosomes (also known as nanochromosomes) that average just 3 kb (Swart *et al.* 2013), each containing its own set of telomeres, regulatory elements, and between 1-8 genes.

Any segment of micronuclear information that is retained in the MAC is called a **macronuclear destined sequence** (abbr. MDS), while any deleted segment that interrupts two MDSs in the same locus is called an **internally eliminated sequence** (abbr. IES). MDSs fuse together by recombination at short (2-20) repeats, called **pointers**, at the ends of MDSs, retaining one copy of the pointer in the MAC. The accuracy of this process is guided by a suite of long and small RNAs ((Nowacki *et al.* 2008; Fang *et al.* 2012), reviewed in Bracht *et al.* (2013) and Yerlici and Landweber (2014)).

Rearrangements can be classified as simple or complex, depending on whether the order and orientation in the macronuclear product is maintained, relative to the germline precursor. Simple rearrangements only involve loss of IESs, and do not require alteration to the order or orientation, relative to the precursor locus. The simplest case is a nanochromosome that originates from a single MDS — i.e., an IES-less locus (Chen *et al.* 2014).

Scrambled rearrangements involve MDSs in different order and/or orientation. Further, we can establish varying levels of complexity, by classifying and ranking rearrangements that require multiple, sequential topological operations to produce the end product, as studied previously by Burns *et al.* (2016a).

The scrambled nature of the MIC permits cases in which MDSs for multiple nanochromosomes can interleave with each other (Chen *et al.* 2014). A special case arises when all MDSs for a MAC chromosome are fully contained between two MDSs (i.e. within an IES) of another nanochromosome, giving rise to nested loci. Interleaving and nesting can also have varying degrees of complexity, or layers of depth, akin to Russian dolls. We note that nested and even Russian doll genes do exist in metazoa, usually as whole genes within introns (Assis *et al.* 2008). Most arise from gene duplication and insertion of young genes or transposons into long introns (Sheppard *et al.* 2016; Wei *et al.* 2013; Gao *et al.* 2012).

In this study, we follow up on previous analyses in *Burns et al.* (2016a), that described the most commonly recurring scrambled patterns across the genome. Here, we present the most elaborate cases of genome rearrangement in *O. trifallax*, which highlight the extraordinary degree of topological complexity that can arise from such a highly plastic genomic architecture.

## METHODS

### Genome sequences from *Oxytricha trifallax*

MIC and MAC sequences from *Oxytricha trifallax* were obtained from the *<mds_ies_db>* website (Burns *et al.* 2016b), in the form of annotated tables, and processed as described in *Burns et al.* (2016a), available from http://knot.math.usf.edu/data/scrambled_patterns/processed_annotation_of_oxy_tri.gff. These were parsed to produce the various types of graphical representations described below using a combination of in-house Python and SQL scripts.

### Graphical representations of scrambled loci

These representations highlight different aspects of chromosomal rearrangement topologies. For the analysis of nested chromosomes we filtered the data to consider only cases with scrambled or non-scrambled MDSs for multiple MAC contigs (nanochromosomes) that derive from a single MIC region. Fig. 1a shows a schematic view of the MIC (precursor) and MAC (product) versions of such a locus, and the correspondence between MDSs and pointers. In addition, we include a condensed linear sequence representation of the MDSs (Fig. 1b), a self-intersecting line corresponding to the MIC region that indicates the topological orientation of the product in a single trace (Fig. 1c), and a chord diagram representation of the pointer list for one of the MAC chromosomes (Fig. 1d).

**Figure 1.**
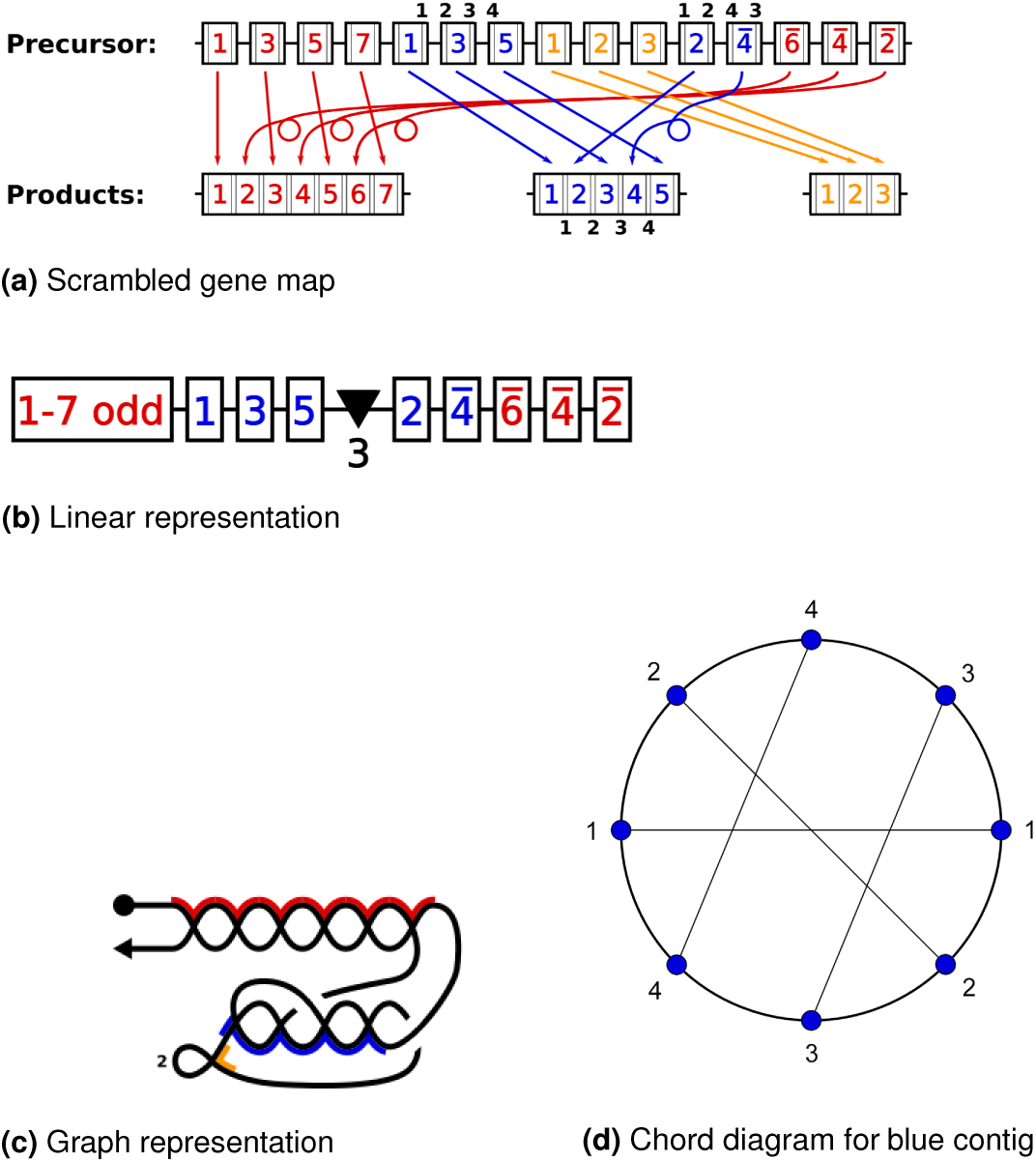
Graphical representation of DNA rearrangements and their topologies. (a) A schematic idealized MIC contig containing MDSs for three MAC contigs, each indicated by a different color. The numbers indicate the final MDS order in the corresponding MAC contig. The numbers with bars above them indicate MDSs in a reverse orientation in the MIC contig relative to the MAC contig. The black numbers in smaller print over the blue MDSs label the pointers for the blue contig. (b) A condensed, linear description of the precursor MIC contig in (a). The notation “1 − 7 odd” represents MDS 1, 3, 5, 7, and the black triangle indicates the presence of three nonscrambled MDSs for the orange MAC contig shown in (a). (c) A graph representing the precursor MIC contig in (a) where the vertices (intersection points) indicate the recom-bination junctions (pointers). The arrowhead indicates the orientation of the precursor MIC contig, reading left to right as shown in (a). The segments highlighted in 3 colors indicate the MAC contigs obtained after joining MDS segments. The uncolored segments correspond to IESs. The vertices of the blue portion correspond to the pointers 1, 2, 3, 4 that join respectively flanking MDSs. The orange portion with a loop corresponds to the middle orange MAC locus with three non-scrambled MDSs. The number 2 indicates removal of two conventional IESs. (d) A chord diagram representing a cyclic arrangement of pointers of MDSs from the blue MAC contig in (a) within its precursor MIC contig. Vertices are labeled in order of pointer appearances in the MIC contig, and chords (line segments within the circle) connect the two copies of the matching pointers.

### Nested chromosomes

We define a locus as nested if all or some of the MDSs for one nanochromosome are surrounded in the MIC by MDSs for a different nanochromosome. Given that nesting can be layered, we define an *interleaving depth index* (IDI) that recursively counts nesting events between the MDSs of a given nanochromosome. For example, in Figures 1 and 2 the IDI values of the orange, blue and red contigs are 0, 1, and 2, respectively. MDSs that map to distinct MAC contigs whose terminal sequences overlap (Chen *et al.* 2014) were not considered in our analysis. Conversely, the *embedding index* (EI) represents the maximal depth of an MDS that resides between the MDSs for another MAC chromosome, counting the layers or levels surrounding it. In Figures 1 and 2, the EI value for each MDS in the red contig is 0, for the blue contig is 1 and for the orange contig is 2. Note that a single MDS can be more deeply embedded than an entire gene locus.

**Figure 2.**
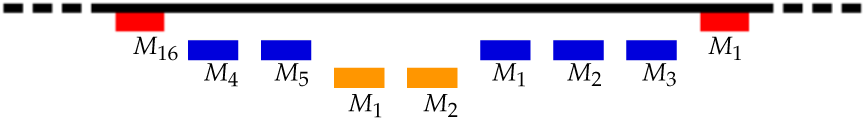
A specific example of three layers of nesting. The two MIC loci for MAC contigs Contig9583.0 (blue) and Contig6683.0 (orange) are nested between Contig6331.0 (red) on the micronuclear contig ctg7180000067077. Not drawn to scale. The red locus has an IDI of 2, the blue locus 1, and the orange locus 0.

### Highly scrambled loci

We previously reported that most scrambled loci in *O. trifallax* contain various combinations of recurrent scrambled patterns, and that recurrent patterns can account for a large majority (96%) of all scrambled genes (Burns *et al.* 2016a). For example, the MAC contig represented by red MDSs in Figure 1 is scrambled, with the MDS order in the MIC locus 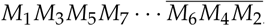, a cluster of odd MDSs in consecutive order, separated from the corresponding even numbered segments in reverse consecutive order. Scrambled genes with an odd-even pattern were first discovered for a single gene in Mitcham *et al.* (1992).

Iterating such recurrent patterns (inversions, translocations, and odd/even splittings) permits a reduction of complexity for each locus (possibly mimicking the evolutionary or developmental steps that occur in nature). Mathematically, we use double occurrence words (defined next) and chord diagrams to describe genome rearrangements.

The order of pointer sequences in a rearrangement map can be used as words, and because every pointer appears twice in the MIC locus, the list of pointers forms a *double occurrence word* (DOW). In the simplest case, pointers that flank a nonscrambled IES between two consecutive MDSs, such as all MDSs in the orange locus, appear in DOWs as pairs of identical, consecutive symbols. Hence the central orange locus in Figure 1, *M*_1_*M*_2_*M*_3_, would be represented by the DOW 1122, where 1 represents the pointer sequence flanking the first IES between *M*_1_and *M*_2_, and 2 is the pointer flanking the second IES, joining *M*_2_to *M*_3_. More generally, pointer *i* is the short repeat (microhomology) present at the end of *M*_*i*_ and beginning of *M*_*i*_+1. Because this study of more complex loci focuses on scrambled patterns, all pairs of neighboring identical pointers can be ignored, and we have eliminated them for simplification. This may also reflect the rearrangement steps during development, since Möllenbeck *et al.* (2008) observed simple IES elimination before MDS reordering or inversion.

For a scrambled example, in Fig. 1(a) and (b), with MDSs labeled as numbers in boxes, “1 − 7 odd” in (b) corresponds to red MDSs *M*_1_*M*_3_*M*_5_*M*_7_, whose corresponding pointer list is 123456. The remaining red boxes appear as numbered 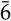, 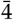, 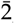, which has a pointer list 654321. Therefore, the pointer list corresponding to the red locus is 123456654321 in the MIC contig, a canonical odd-even pattern (and the 66 in the center is a scrambled pointer junction). Usually, in the *Oxytricha* genome such odd-even patterns appear in the DOW as segments that are repeated or reversed. For example, 1234 … 1234 is a *repeat word* corresponding to *M*_1_*M*_3_*M*_5_…*M*_2_*M*_4_ and 1234 … 4321 is a *return word* corresponding to MDS sequence 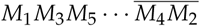

The pointer numbers at the top of the scrambled blue MDSs in Figure 1a read 12341243. Thus the DOW representing this MAC contig is 12341243. In this case 12 ⋅ 12 is a *maximal repeat word* inside. The first step of reduction removes these maximal repeat words. The remaining word 3443 is a *return word* or perfect inverted repeat. In a second step of reduction, we eliminate this return word, leaving the empty word *ε*. Note that we use this iterative process to characterize the complexity of a scrambled locus, but it may or may not reflect either the pathway for gene descrambling or the evolutionary steps that led to its scrambling.

We describe these patterns by chord diagrams associated with each DOW. In a chord diagram the symbols of a DOW are placed in order from a reference point on a circle (marked by a small bar). The diagram is obtained by connecting two identical symbols placed on a circle with a chord (see Fig. 1d). The chord diagrams corresponding to repeat and return words have the form depicted in Fig. 3.

**Figure 3.**
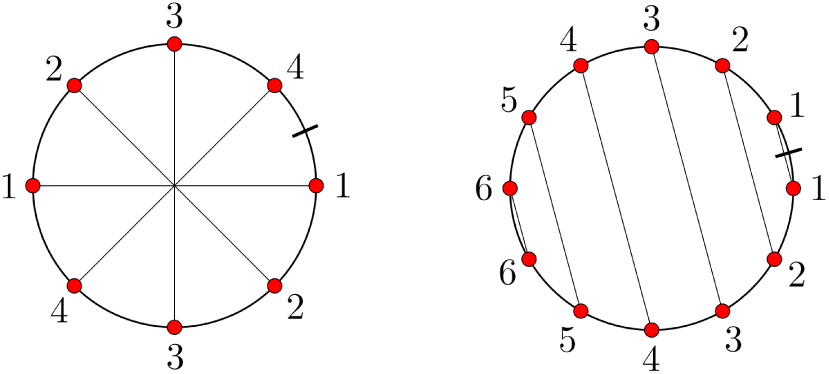
The chord diagrams associated with the repeat word 12341234 (left) and the return word 123456654321 (right).

**Figure 4.**
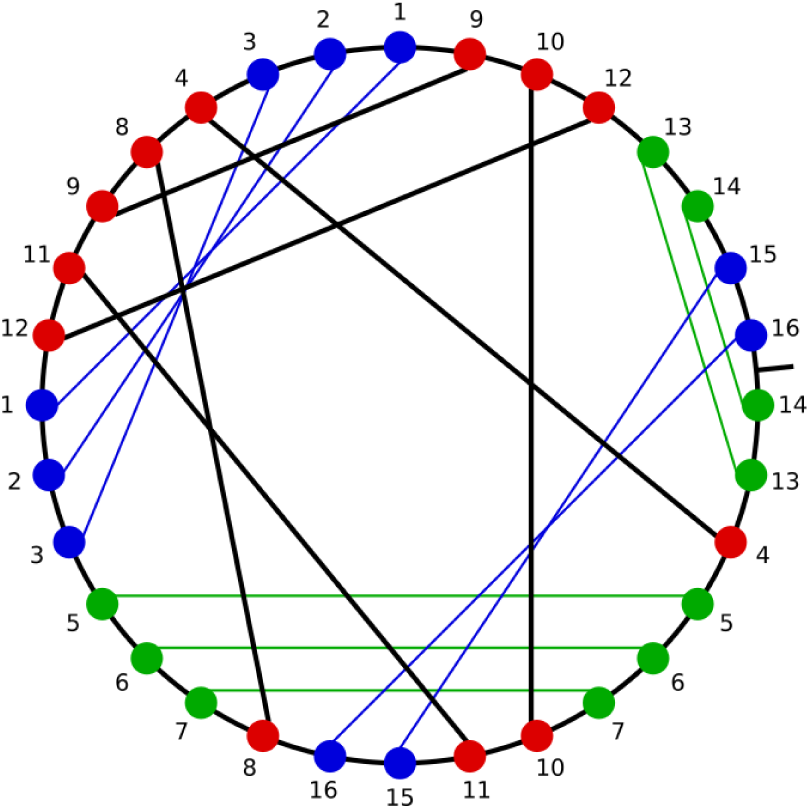
The chord diagram associated with Contig582.1. Repeat/return patterns are highlighted in blue and green, respectively. Red nodes are not found in repeat or return words.

Repeat and return words can appear nested within one another in *Oxytricha*’s scrambled genome. For example, the word 121*342***5665***34* contains the return word **5665** nested within the repeat word *3434*. After removal of both 5665 and 3434, the word reduces to 1212 which is again a repeat word, which further reduces to *ε*. In *Burns et al.* (2016a) we found that over 90% of scrambled contigs can be reduced by various combinations of these two operations. Therefore their topological complexity can be broken down into simple steps, which may have arisen by layer upon layer of germline translocations that introduce or propagate odd-even patterns (Chang *et al.* 2005; Landweber 1998).

Here, we identify rare patterns in the genome and exceptional cases of nested genes. We do so by recursively removing repeat and return patterns for the genome-wide dataset analyzed in *Burns et al.* (2016a), and we retain those contigs whose descriptions cannot be further reduced.

## RESULTS AND DISCUSSION

### The *Oxytricha trifallax* genome contains deeply nested loci

The interleaving depth index (IDI) was computed for 15,811 MAC contigs and is summarized in Table 1. Two exceptional MIC loci each contain the nested precursor segments for four other MAC contigs, and this represents the highest level of nesting, IDI=4 (or 5 nested genes). These two MIC loci are shown in Figures 5a and 5b and their topological representations are shown in Figures 6a and 6b, respectively.

**Table 1.**
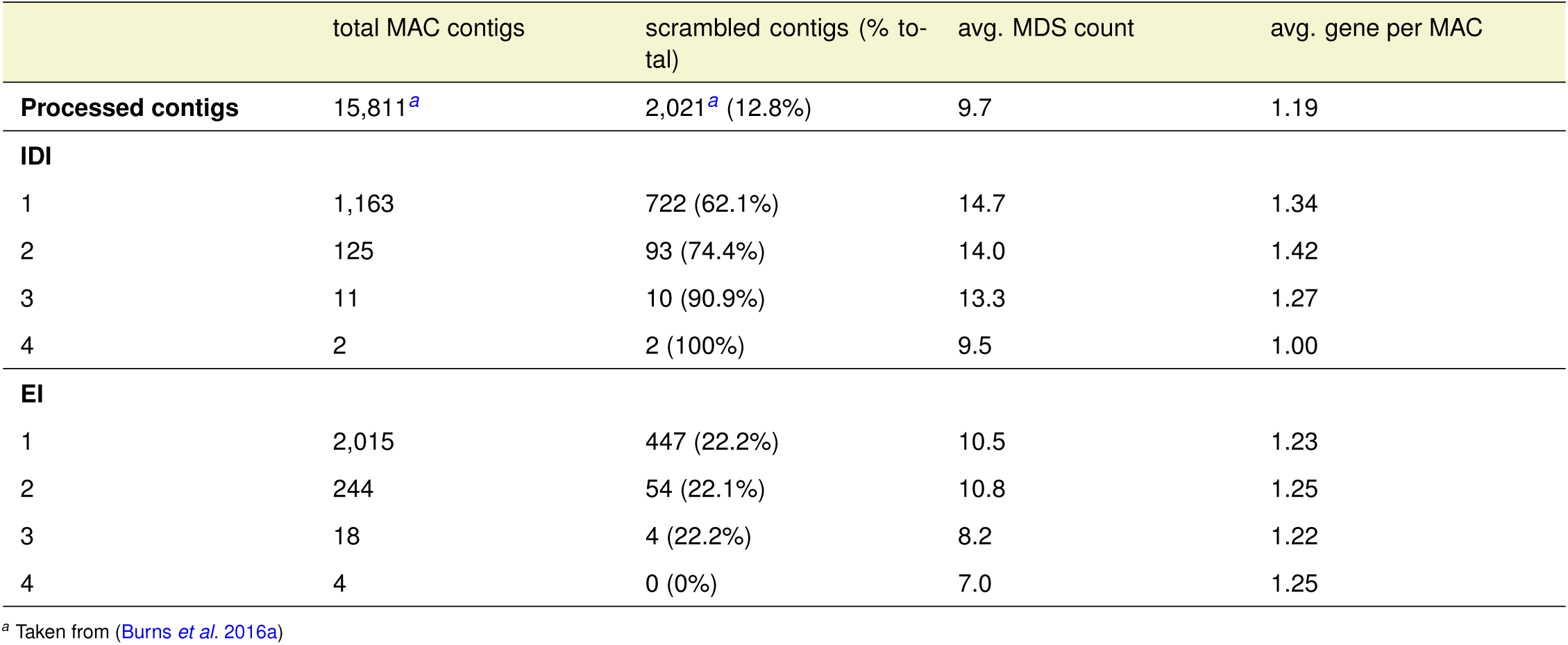
Nested and embedded germline loci

|  | total MAC contigs | scrambled contigs (% total) | avg. MDS count | avg. gene per MAC |
| --- | --- | --- | --- | --- |
| **Processed contigs** | 15,811<sup>a</sup> | 2,021<sup>a</sup> (12.8%) | 9.7 | 1.19 |
| **IDI** |  |  |  |  |
| 1 | 1,163 | 722 (62.1%) | 14.7 | 1.34 |
| 2 | 125 | 93 (74.4%) | 14.0 | 1.42 |
| 3 | 11 | 10 (90.9%) | 13.3 | 1.27 |
| 4 | 2 | 2 (100%) | 9.5 | 1.00 |
| **EI** |  |  |  |  |
| 1 | 2,015 | 447 (22.2%) | 10.5 | 1.23 |
| 2 | 244 | 54 (22.1%) | 10.8 | 1.25 |
| 3 | 18 | 4 (22.2%) | 8.2 | 1.22 |
| 4 | 4 | 0 (0%) | 7.0 | 1.25 |

<sup>a</sup> Taken from (Burns *et al.* 2016a)

**Figure 5.**
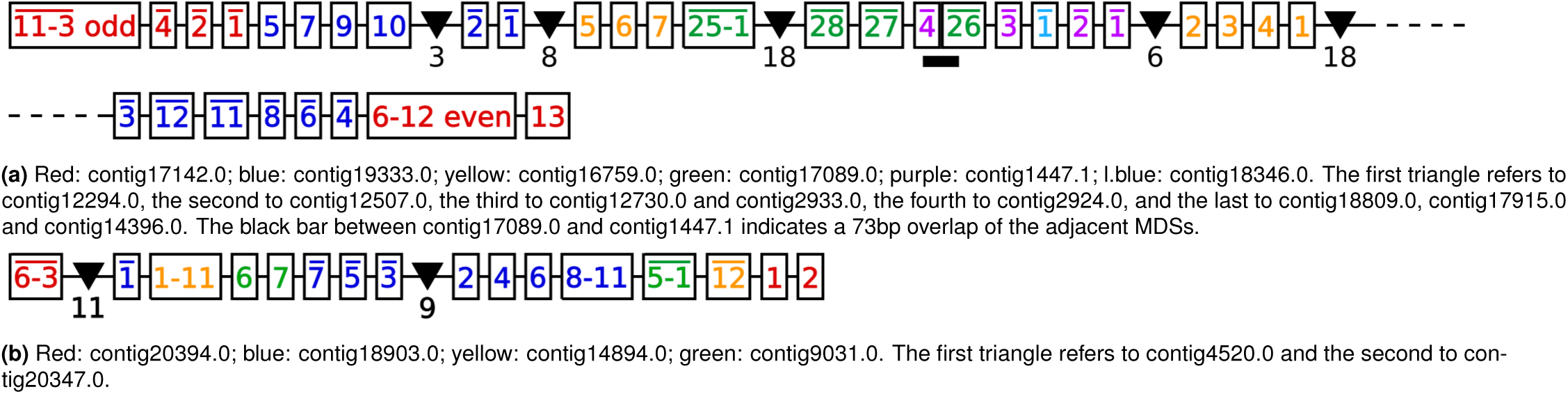
Germline maps for two MIC contigs, the "Russian Doll" loci (a) ctg7180000087484 and (b) ctg7180000069233 containing several nested genes. The numbers indicate the positions of the MDSs in the sequence along the MAC contig. Inverted MDSs are marked with bars above the numbers. Multiple sequential MDSs are condensed with a dash and the words ‘even’ or ‘odd’ to indicate if only even or odd numbered MDSs, respectively, are present in that string. Triangles denote the presence of additional nonscrambled MAC loci with the number below each triangle indicating the number of nonscrambled MDSs for that locus.

**Figure 6.**
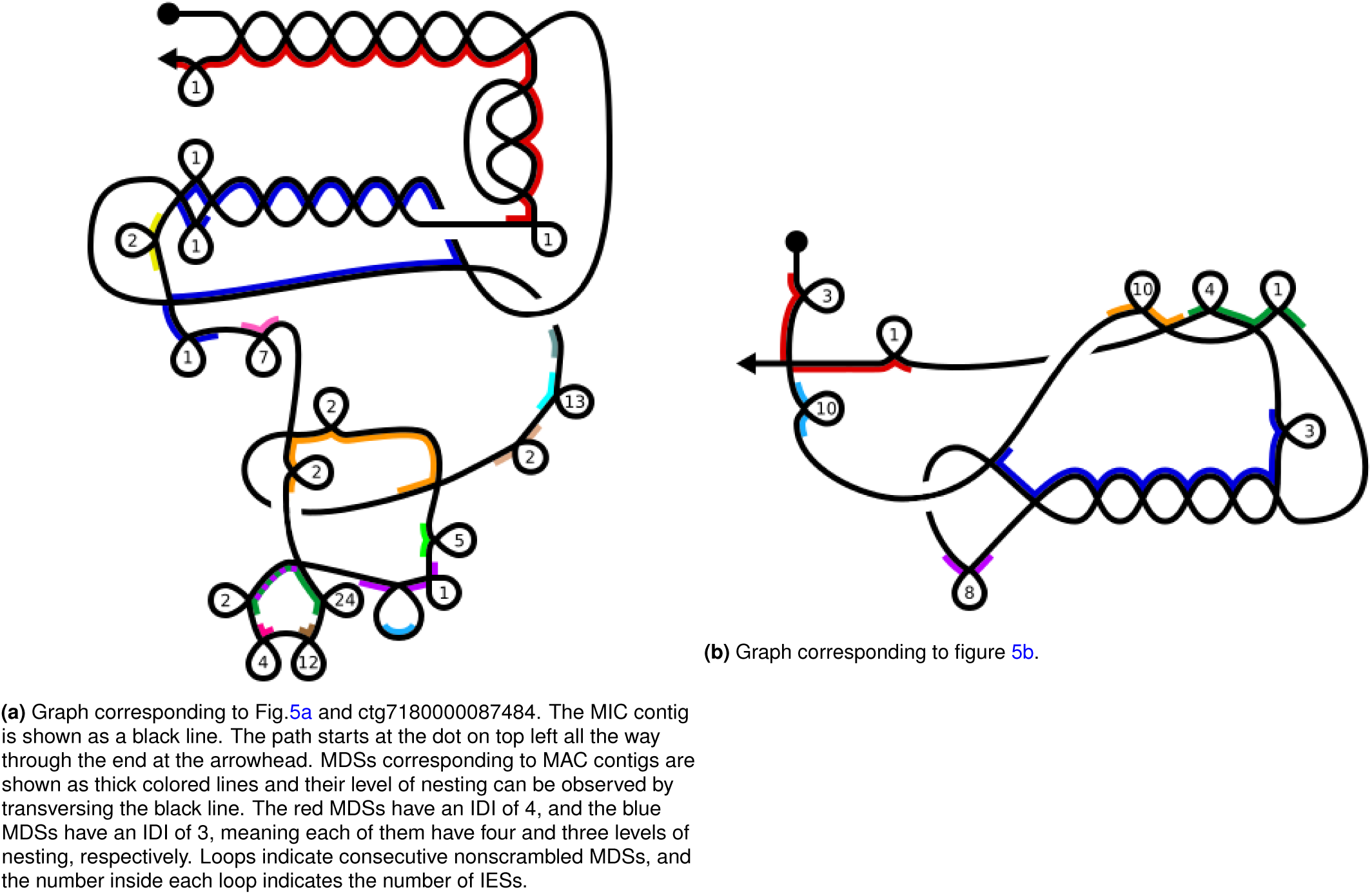
Graphical representations of the recombinations of the MIC loci depicted in Figure 5. Not drawn to scale. The vertices indicate the recombination sites corresponding to alignments of pointers. Loops mark conventional (nonscrambled) IESs, which occur whenever sequential MDSs are present successively on the MIC. The number inside each loop indicates the number of conventional IESs. Colored edges indicate MDSs of MAC contigs. The colors of the MAC contigs are the same as in Figure 5. More colors were added to indicate the nonscrambled contigs corresponding to the triangles in Figure 5.

1301 (8%) of MAC contigs contain MDSs for other nanochromosomes nested within them in the MIC (IDI of 1 or greater). Among these, scrambling is more common, when compared with the genome-wide rate of scrambling (*χ*^2^ test, p<0.01). Furthermore, higher IDI values correlate with a higher proportion of scrambled loci (Spearman’s *ρ*=1, p<0.05). EI, on the other hand, does not correlate with the presence or absence of scrambling. 22% of MAC chromosomes are scrambled in the MIC for all EI values greater than zero (Table 1), which is indistinguishable from the genome-wide average (Chen *et al.* 2014). There is no significant correlation between the number of MDSs per MAC locus and IDI (or EI), but among nested loci, we note that those within inner layers tend to have fewer MDSs than those in outer layers, perhaps as a result of recent insertion.

The evolutionary steps that produce nested genes may favor nanochromosomes that are themselves scrambled, since their longer pointers (Chen *et al.* 2014) might facilitate rearrangement across greater distances. Additionally, the distance between two MDSs in the MIC that are consecutive in the MAC is typically longer for scrambled vs. nonscrambled cases (Chen *et al.* 2014), which provides more opportunity for insertion of other MDSs between them. The elements they contain (the innermost Russian dolls), themselves, do not display any bias for being scrambled or nonscrambled.

We further classified loci as interleaving or embedded. Interleaving occurs when some (but not all) MDSs of a nanochromosome reside between MDSs for another MAC contig. A locus is considered embedded when all MDSs for a MAC contig are contained within the MIC locus of another MAC contig (Table 2). Interleaved cases show the highest proportion of scrambled loci.

**Table 2.**
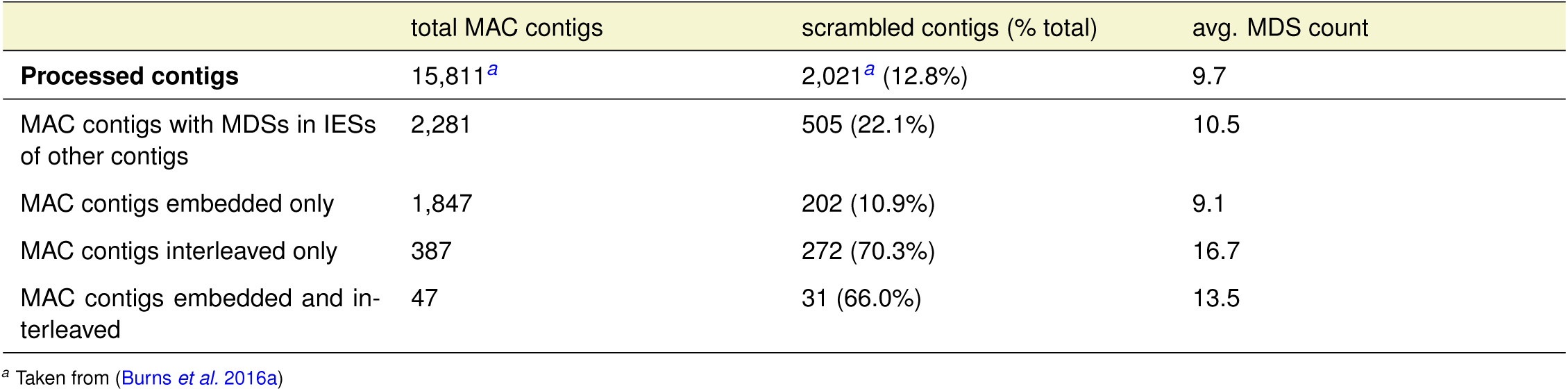
Embedding and Interleaving. The number of MAC contigs which have MDSs with IESs of other MAC contigs, as well as the numbers of the subsets of those MAC contigs that are only found embedded^1^ in other contigs, vs. only interleaved^2^, and those that can be found embedded in some and interleaved with other MAC contigs. ^1^: A MAC contig is embedded in another if all its MDSs are found on a single IES of the other MAC contig. To say that a MAC contig can only be found embedded in others, means that whenever one of its MDSs is found within an IES of another MAC contig, all of the MDSs reside within the same IES. ^2^: A MAC contig is interleaved with another if at least one MDS of each contig is found on an IES of the others. To say that a MAC contig can only be found interleaved with others, means that whenever an MDS of it is found on another MAC contigs, that contig also has an MDS on the IES of the original contig.

|  | total MAC contigs | scrambled contigs (% total) | avg. MDS count |
| --- | --- | --- | --- |
| **Processed contigs** | 15,811<sup>a</sup> | 2,021<sup>a</sup> (12.8%) | 9.7 |
| MAC contigs with MDSs in IESs of other contigs | 2,281 | 505 (22.1%) | 10.5 |
| MAC contigs embedded only | 1,847 | 202 (10.9%) | 9.1 |
| MAC contigs interleaved only | 387 | 272 (70.3%) | 16.7 |
| MAC contigs embedded and interleaved | 47 | 31 (66.0%) | 13.5 |

<sup>a</sup> Taken from (Burns *et al.* 2016a)

### Highly scrambled and atypical loci

The rearrangement topology of 96% of the scrambled chromosomes in *O. trifallax*, even large ones with hundreds of scrambled MDSs (Chen *et al.* 2014), can be explained by combinations of two types of operations: the mixture of odd-even patterns, or repeat and return words (Burns *et al.* 2016a). However, 176 scrambled loci were identified that could not be reduced by iterative application of these reduction operations. In this section we describe the 22 cases that contain at least 4 scrambled pointers after this reduction.

Figure 4 shows an example of one the 22 contigs, with nested pairs of repeat and return words in blue and green, respectively. The reduction process leaves 6 scrambled pointers with DOW 4, 10, 11, 8, 12, 11, 9, 8, 4, 9, 10, 12 (commas added to separate the symbols). This word has no appearances of a repeat or return pattern.

Table 3 lists the predicted genes encoded on these 22 nanochromosomes (Burns *et al.* 2016a), and Supplemental Material S1 describes the MIC loci, the MDSs for these scrambled cases, and their scrambled rearrangement paths as chord diagrams. All but 7 chromosomes encode single genes. The remaining 7 chromosomes each encode two or three genes. Scrambled pointers that cannot be explained as recursive combinations of odd-even patterns (considered irreducible in Burns *et al.* (2016a)) are shown as dark lines in the chord diagrams (and red nodes in Figure 4).

**Table 3.**
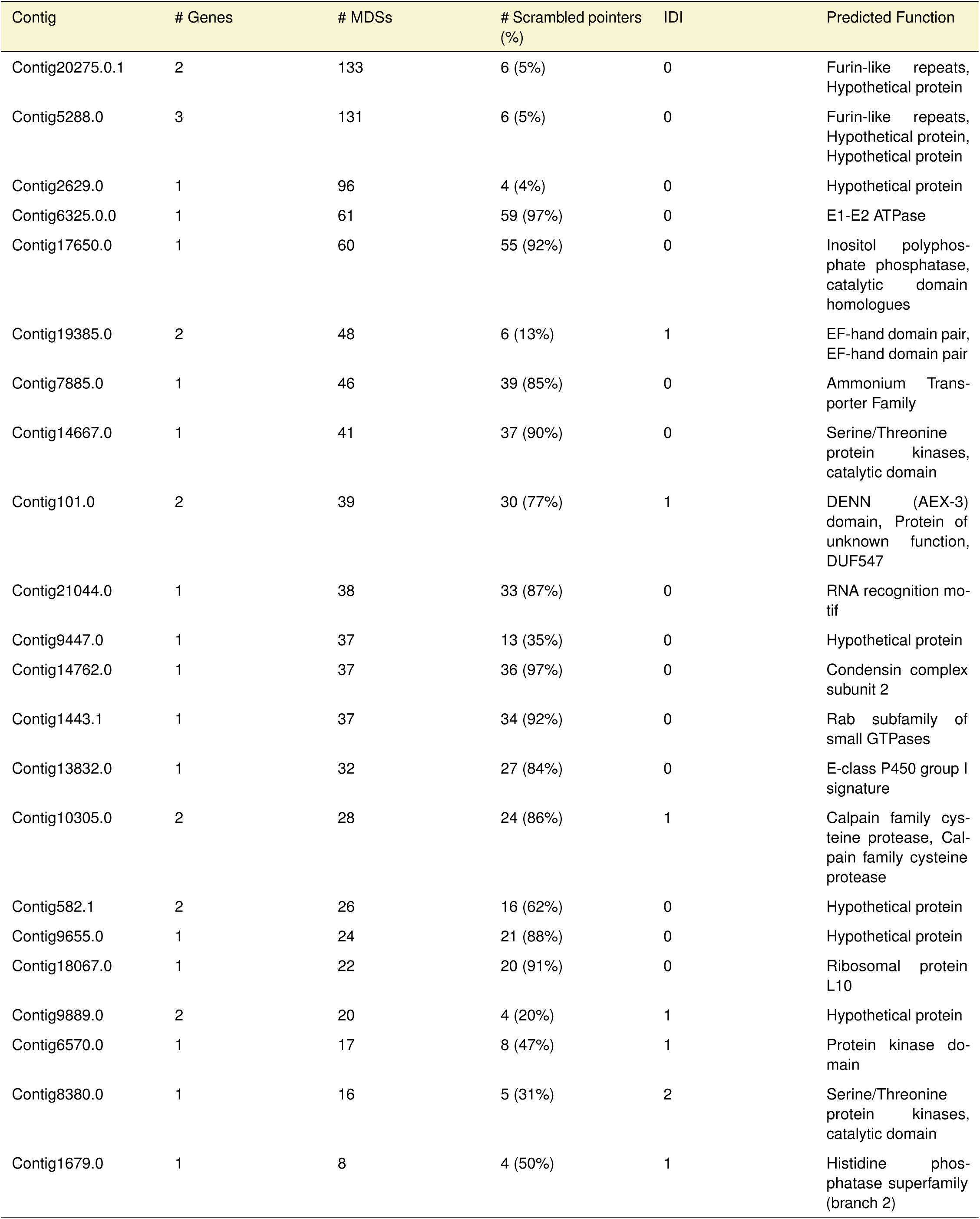
Highly Scrambled and Atypical MAC Chromosomes

Some of the most complex cases have over 50 scrambled MDSs. The two largest cases are over 100 MDSs; however, nearly all of those MDSs are nonscrambled. Only seven MDSs are scrambled in these loci and they share the same chord diagrams, but these two MAC chromosomes are actually paralogous (75% similar at the nucleotide level), arising from a tandem duplication of a 77 kb region in the MIC genome.

The simplest case among the 22 loci is just 8 MDSs in the order 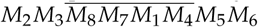. After eliminating t<u>he 3 conv</u>entional (nonscrambled) IESs, this map simplifies to 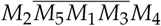, which requires descrambling at just 4 pointer junctions and contains no ostensible pattern. However, its scrambled pointer list 12413234 contains the pattern 121323 (as a substring after pointer 4 is recombined or removed).

The graph representation of this pattern is a tangled cord, which contains slinky-like coiled circles (Burns *et al.* 2013). The simplest form of an MDS rearrangement map corresponding to a tangled cord is 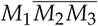. Its pointer list 1212 is also a repeat word, however, that equally describes the odd-even map *M*_1_*M*_3_*M*_2_, as well as 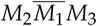, which is not odd-even because of the inversion. Therefore, a *tangled cord* is defined recursively by the DOW, or pointer list, that corresponds to the MDS rearrangement map: 1212 is the simplest case. The next is 121**3**2**3**, and 12132**4**3**4** etc. The DOW for the pointer list grows by adding a new pair of identical symbols after the last and penultimate symbols of the previous DOW.

For another example, the tangled cord pointer list 121323 describes both of the following rearrangement maps, which appear among the other 154 MIC contigs with fewer than 4 scrambled pointers remaining after eliminating all odd-even layers: 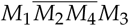 and *M*_2_*M*_1_*M*_4_*M*_3_ in the MIC loci for MAC contigs 567.1 and contig 17193.0, respectively.

We find that the tangled cord can be detected in all but one (contig101.0) of the 22 loci in this study. This includes the cases with the fewest scrambled MDSs (Table 3 and Supplement S1). Such embedding of the tangled cord appears highly prevalent among these 22 loci, but whether the pattern appears more often than would be expected by chance remains to be tested (Jonoska *et al.* 2017).

## PERSPECTIVES

Our results show that a great diversity of scrambled gene maps are present in the germline genome of *O. trifallax*. The presence of such highly nested architectures was a surprise and suggests that layers upon layers of MDS and gene translocations constantly alter the genome, while the detection of highly scrambled patterns reveals architectures that go beyond simple twists and turns of the DNA. Several open questions remain: how do these patterns arise in evolution? Do nested and highly scrambled patterns accumulate gradually or in a catastrophic event, like chromothripsis (Maher and Wilson 2012)? Are the combinations of patterns serendipitous, or is there an biological process that drives the introduction of higher levels of scrambling? Future studies should address population variation, and measure the level of variation, in chromosome structures at different scales of evolutionary divergence. In particular, surveys of the orthologous loci for the notable cases studied here in earlier diverged spirotrichous ciliates (as in Chang *et al.* (2005) and Hogan *et al.* (2001)) have the potential to reveal much about the evolutionary steps that gave rise to such complex, intertwined genome architectures. Furthermore, the current study offers new metrics of topological genome complexity, that go beyond the linear nature of eukaryotic chromosomes and consider their deeply structured and layered history.

## LITERATURE CITED

Assis, R., A. S. Kondrashov, E. V. Koonin, and F. A. Kondrashov, 2008 Nested genes and increasing organizational complexity of metazoan genomes. Trends Genet. 24: 475–478.

Bracht, J. R., W. Fang, A. D. Goldman, E. Dolzhenko, E. M. Stein, and L. F. Landweber, 2013 Genomes on the edge: programmed genome instability in ciliates. Cell 152: 406–416.

Burns, J., E. Dolzhenko, N. Jonoska, T. Muche, and M. Saito, 2013 Four-regular graphs with rigid vertices associated to DNA recombination. Discrete Applied Mathematics 161: 1378–1394.

Burns, J., D. Kukushkin, X. Chen, L. F. Landweber, M. Saito, and N. Jonoska, 2016a Recurring patterns among scrambled genes in the encrypted genome of the ciliate Oxytricha trifallax. Journal of Theoretical Biology 410: 171–180.

Burns, J., D. Kukushkin, K. Lindblad, X. Chen, N. Jonoska, and L. F. Landweber, 2016b : a database of ciliate genome rearrangements. Nucleic Acids Research 44: D703–D709, Avail- able at: http://knot.math.usf.edu/mds_ies_db/index.html, accessed October 2017.

Chang, W. J., P. D. Bryson, H. Liang, M. K. Shin, and L. F. Landweber, 2005 The evolutionary origin of a complex scrambled gene. Proc. Natl. Acad. Sci. U.S.A. 102: 15149–15154.

Chen, X., J. R. Bracht, A. D. Goldman, E. Dolzhenko, D. M. Clay, E. C. Swart, D. H. Perlman, T. G. Doak, A. Stuart, C. T. Amemiya, R. P. Sebra, and L. F. Landweber, 2014 The architecture of a scrambled genome reveals massive levels of genomic rearrangement during development. Cell 158: 1187–1198.

Fang, W., X. Wang, J. R. Bracht, M. Nowacki, and L. F. Landweber, 2012 Piwi-interacting RNAs protect DNA against loss during Oxytricha genome rearrangement. Cell 151: 1243–1255.

Gao, C., M. Xiao, X. Ren, A. Hayward, J. Yin, L. Wu, D. Fu, and J. Li, 2012 Characterization and functional annotation of nested transposable elements in eukaryotic genomes. Genomics 100: 222–230.

Hogan, D. J., E. A. Hewitt, K. E. Orr, D. M. Prescott, and K. M. Muller, 2001 Evolution of IESs and scrambling in the actin I gene in hypotrichous ciliates. Proc. Natl. Acad. Sci. U.S.A. 98: 15101–15106.

Jonoska, N., L. Nabergall, and M. Saito, 2017 Patterns and distances in words related to DNA rearrangement. Fundamenta Informaticae 154: 225–238.

Landweber, L. F., 1998 The Evolution of DNA Computing: Nature's Solution to a Path Problem. IEEE Proceedings of Symposia on Intelligence and Systems 98 pp. 133–139.

Maher, C. A. and R. K. Wilson, 2012 Chromothripsis and human disease: piecing together the shattering process. Cell 148: 29–32.

Mitcham, J. L., A. J. Lynn, and D. M. Prescott, 1992 Analysis of a scrambled gene: the gene encoding alpha-telomere-binding protein in Oxytricha nova. Genes Dev. 6: 788–800.

Möllenbeck, M., Y. Zhou, A. R. Cavalcanti, F. Jönsson, B. P. Higgins, W.-J. Chang, S. Juranek, T. G. Doak, G. Rozenberg, H. J. Lipps, and L. F. Landweber, 2008 The pathway to detangle a scrambled gene. PloS one 3: e2330.

Nowacki, M., V. Vijayan, Y. Zhou, K. Schotanus, T. G. Doak, and L. F. Landweber, 2008 RNA-mediated epigenetic programming of a genome-rearrangement pathway. Nature 451: 153–158.

Prescott, D. M., 1994 The DNA of ciliated protozoa. Microbiol. Rev. 58: 233–267.

Sheppard, A. E., N. Stoesser, D. J. Wilson, R. Sebra, A. Kasarskis, L. W. Anson, A. Giess, L. J. Pankhurst, A. Vaughan, C. J. Grim, H. L. Cox, A. J. Yeh, C. D. Sifri, A. S. Walker, T. E. Peto, D. W. Crook, and A. J. Mathers, 2016 Nested Russian Doll-Like Genetic Mobility Drives Rapid Dissemination of the Carbapenem Resistance Gene blaKPC. Antimicrob. Agents Chemother. 60: 3767–3778.

Swart, E. C., J. R. Bracht, V. Magrini, P. Minx, X. Chen, Y. Zhou, J. S. Khurana, A. D. Goldman, M. Nowacki, K. Schotanus, S. Jung, R. S. Fulton, A. Ly, S. McGrath, K. Haub, J. L. Wiggins, D. Stor- ton, J. C. Matese, L. Parson, W.-J. Chang, M. S. Bowen, N. A. Stover, T. A. Jones, S. R. Eddy, G. A. Herrick, T. G. Doak, R. K. Wilson, E. R. Mardis, and L. F. Landweber, 2013 The Oxytricha trifallax macronuclear genome: A complex eukaryotic genome with 16,000 tiny chromosomes. PLoS Biology 11.

Wei, L., M. Xiao, Z. An, B. Ma, A. S. Mason, W. Qian, J. Li, and D. Fu, 2013 New insights into nested long terminal repeat retro-transposons in Brassica species. Mol Plant 6: 470–482.

Yerlici, V. T. and L. F. Landweber, 2014 Programmed Genome Rearrangements in the Ciliate Oxytricha. Microbiol Spectr 2.

